# Preferential binding of ADP-bound mitochondrial HSP70 to the nucleotide exchange factor GRPEL1 over GRPEL2

**DOI:** 10.1101/2024.05.16.594508

**Authors:** Pooja Manjunath, Gorazd Stojkovič, Svetlana Konovalova, Sjoerd Wanrooij, Kristian Koski, Henna Tyynismaa

## Abstract

Human nucleotide exchange factors GRPEL1 and GRPEL2 play pivotal roles in the ADP-ATP exchange within the protein folding cycle of mitochondrial HSP70 (mtHSP70), a crucial chaperone facilitating protein import into the mitochondrial matrix. Studies in human cells and mice have indicated that while GRPEL1 serves as an essential co-chaperone for mtHSP70, GRPEL2 has a role regulated by stress. However, the precise structural and biochemical mechanisms underlying the distinct functions of the GRPEL proteins have remained elusive. In our study, we present evidence revealing that ADP-bound mtHSP70 exhibits remarkably higher affinity for GRPEL1 compared to GRPEL2, with the latter experiencing a notable decrease in affinity upon ADP binding. Utilizing Alphafold modelling, we propose that the interaction between GRPEL1 and mtHSP70 can induce the opening of the nucleotide binding cleft of the chaperone, thereby facilitating the release of ADP, whereas GRPEL2 lacks this capability. This study elucidates how ADP binding to mtHSP70 enhances its affinity for GRPEL1, contrasting with its interaction with GRPEL2. Additionally, our findings suggest that the redox-regulated Cys87 residue in GRPEL2 does not play a role in dimerization but rather reduces its affinity for mtHSP70. Our findings on the structural and functional disparities between GRPEL1 and GRPEL2 may have implications for mitochondrial protein folding and import processes under varying cellular conditions.

## 1 INTRODUCTION

Mitochondria are major metabolic regulator organelles that import most of their proteome from the cytosol. The molecular chaperone mtHSP70 of the (heat shock protein) HSP70 family plays a specialised role in the translocation of preproteins into the mitochondrial matrix and in their proper folding (Bracher & Verghese, 2023; Kang et al, 1990). Its protein binding/release cycle is regulated by co-chaperones, which for the bacterial Hsp70 homologue DnaK were identified as DnaJ and the nucleotide exchange factor (NEF) GrpE (Hosfelt et al, 2022; Liberek et al, 1991). ATP hydrolysis by DnaJ stabilises the interaction of substrate protein with DnaK, whereas GrpE triggers ADP to ATP exchange, accelerating substate release, which is rate-limiting in the cycle (Szabo et al, 1994; Xiao et al, 2024). Prokaryotic and eukaryotic Hsp70 systems are functionally similar; however, in eukaryotes, a GrpE-like NEF is found only in mitochondria (Harrison, 2003). The most studied mitochondrial NEF is Mge1, the sole GrpE-like protein in *Saccharomyces cerevisiae*, which is essential for viability (Bolliger et al, 1994; Krzewska et al, 2001; Laloraya et al, 1994). Interestingly, two distinct GrpE-like mitochondrial proteins exist in mammals, GRPEL1 and GRPEL2 (Goswami et al, 2010; Naylor et al, 1996; Naylor et al, 1998; Srivastava et al, 2017), but the structural attributes to their function are not well known.

Bacterial GrpE and yeast Mge1 interact with their respective chaperones as dimers (Harrison et al, 1997; Moro & Muga, 2006; Wu et al, 2012). Notably, human GRPEL1 and GRPEL2 are functionally homodimers (Konovalova et al, 2018), but their stoichiometric interaction with mtHSP70 is yet to be addressed. High-resolution structural information is available only for bacterial GrpE (Harrison et al., 1997; Nakamura et al, 2010; Wu et al., 2012; Xiao et al., 2024). Analysis of the GrpE-DnaK complexes from *Escherichia coli* (PDB-ID1DKG) and *Mycobacterium tuberculosis* (PDB-ID 8GB3) revealed a 2:1 stoichiometry, characterized by an asymmetric interaction between one of the monomers of GrpE dimer and DnaK (Harrison et al., 1997; Xiao et al., 2024). Conversely, when structurally analyzed, the *Geobacillus kaustophilus* GrpE homodimer bound to DnaK exhibited a 2:2 stoichiometry (PDB-ID 4ANI), where each GrpE monomer interacted with its corresponding DnaK monomer, leading to a stiffer dimeric GrpE structure (Wu et al., 2012). While interaction diversity exists among NEFs, the primary function is consistent: binding to the ADP-bound nucleotide binding domain (NBD) of the chaperones to facilitate ADP release (Bracher & Verghese, 2023). Recent studies show that the interaction of GrpE with DnaK serves to allosterically regulate the chaperone, leading to release of ADP as well as the folded protein from the substrate binding domain (SBD) (Rossi et al, 2024; Xiao et al., 2024).

GRPEL1 is conserved across all metazoans, whereas GRPEL2 is found in vertebrates, and both have ubiquitous expression in human tissues (Konovalova et al., 2018; Naylor et al., 1998). However, evidence now indicates that GRPEL1, but not GRPEL2, is the essential housekeeping NEF for mtHsp70 in mammalian cells: i) Only human GRPEL1 was able to complement yeast Mge1 (Srivastava et al., 2017), ii) Mitochondrial protein import was not impaired in human GRPEL2 knockout cells (Konovalova et al., 2018), iii) Human variation data indicates that the GRPEL1 gene is not tolerant to loss-of-function variants, unlike GRPEL2 (Konovalova et al., 2018), iv) Mouse knockout of GRPEL1 was lethal in early development, and the depletion of GRPEL1 in skeletal muscles of adult mice led to severe muscle atrophy and premature death, showing that GRPEL2 was not able to compensate for GRPEL1 loss (Neupane et al, 2022). In contrast, we previously reported that GRPEL2 may play a specific role in stress sensing similar to Mge1, which is an oxidative stress sensor in yeast (Marada et al, 2013). We observed that in cultured human cells, the dimerization of GRPEL2 increased under oxidative stress and was dependent on redox-regulated Cys87 (Konovalova et al., 2018). In agreement, a competitive cysteine-reactive profiling study recently identified Cys87 as the most redox-sensitive cysteine of GRPEL2 (Kisty et al, 2023).

In this study, we aimed to explicate the fundamental structural, biochemical, and biophysical differences between GRPEL1 and GRPEL2 to increase our understanding of their potential roles. In our investigation, we revealed an enhanced interaction specifically between ADP-bound mtHSP70 and GRPEL1. Furthermore, we show that disulphides affect the stability of GRPEL2 and impede its interaction with mtHSP70. These findings further strengthen the importance of GRPEL1 as the main NEF for mtHSP70 in human mitochondria.

## 2 RESULTS

### 2.1 Sequence comparison of the GRPELs highlights the unique cystines in GRPEL2

To explore the distinctions in interactions between mtHSP70 and both GRPELs, we examined the amino acid sequence variances among the NEFs and compared them with yeast Mge1 and three bacterial GrpE structures (4ANI, 1DKG and 8GB3) (Figure 1a). We further studied the cysteine pattern and structure of the GRPELs by calculating the dimeric models with the help of AlphaFold 2 (Jumper et al, 2021) (Figure 1b and c). In GRPEL2, Cys87, the most redox-sensitive cysteine (Kisty et al., 2023), locates in the middle of the long α1-helix, and in the dimeric AlphaFold2 model, it indeed forms a disulphide bridge with the corresponding cysteine of the dimer partner (Figure 1c). As seen in the sequence alignment of the GrpE family of proteins (Figure 1a), Cys87 is only found in GRPEL2. Similarly, the next cysteine, Cys97, located in the α1-helix, is only present in GRPEL2. However, Cys97 is not forming a disulphide bridge according to the AlphaFold2 model but points to the bulk solvent. The next two cysteines, Cys110 and Cys126, both located in the α2-helix, are found in GRPEL1 and GRPEL2, but are not conserved in yeast or bacterial homologues (Figure 1a). These two cysteines have also been shown to possess redox-sensitivity properties (Kisty et al., 2023). GRPEL2 has an additional unique cysteine at the end of β2 (Cys182). Both GRPEL1 and GRPEL2 have one N-terminal cysteine (C28 and C44, respectively) in the random-coil region. These sequence comparisons suggest that unique cystines in GRPEL2 may perform specific roles under stress conditions.

**Figure 1.**
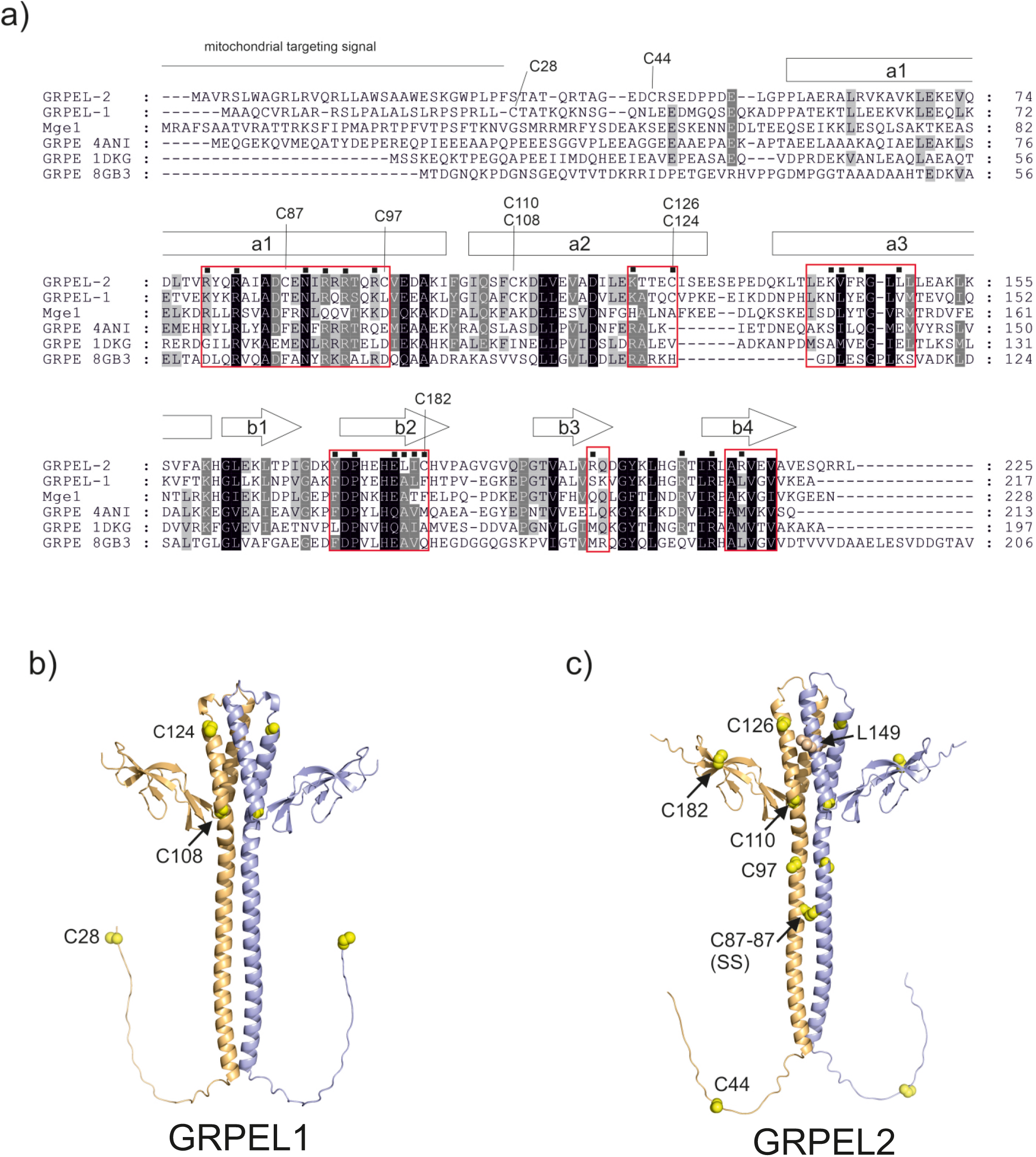
Sequence comparison of the GRPELs highlights the unique cystines in GRPEL2. (a) Sequence alignment of GrpE protein family members including GRPEL1 and GRPEL2, the yeast Mge1, and the three bacterial GrpE proteins from *E. coli*, *G. kaustophilus* and *M. tuberculosis*, which have crystal structures determined (PDB codes 4ANI, 1DKG and 8GB3, respectively). The cysteines in GRPEL2 and GRPEL1 are highlighted and labelled. The secondary structure elements of GRPEL2 are shown above the sequences as predicted by the AlphaFold2 model. The regions that are shown to interact with HSP70 by structural GrpE studies (Harrison et al., 1997) (Wu et al., 2012) and by the modelling studies of this study (by AlphaFold2 Multimer, see Figure S1) are shown with boxes around the sequence regions. Some key residues forming the interactions are highlighted with black squares. Met155 is a critical redox sensor in Mge1 (Marada et al., 2013). The corresponding Leu149 of GRPEL2 is highlighted with an arrow. (b) AlphaFold2 model of the GRPEL1 dimer as calculated using AlphaFold2 Colab (Evans et al., 2021; Jumper et al., 2021). The chains are coloured as light orange and blue. The cysteine residues are shown with yellow spheres. c) AlphaFold2 model of the GRPEL2 dimer. The colouring scheme is the same as in panel b). All the cysteines, and the Leu149, corresponding to Met155 of Mge1, are shown with light brown spheres. GRPEL models exclude the mitochondrial targeting signal in the N-terminus of both chains.

### 2.2 Purified-soluble human GRPELs exist in dimeric form and cysteines have no role in oligomerization

To investigate the biophysical and biochemical differences between GRPELs, we purified recombinant human GRPEL1, GRPEL2, and mtHSP70 as well as mutant variants of GRPEL2 with either a Cys87 to Ala (GRPEL2-C87A) or a Cys97 to Ala (GRPEL2-C97A). The mtHSP70 and GRPEL constructs were purified by affinity chromatography and size-exclusion chromatography (SEC). All the protein obtained was soluble, and the concentrations were typically between 4-8mg/ml. After the purification step, the purified GRPEL proteins resulted in a single protein band with a molecular mass of around 25 kDa in SDS-PAGE while the purified mtHSP70 exhibited greater heterogeneity, with the primary band aligning with a size of around 70 kDa (Figure 2a). However, the gel filtration profile displayed several peaks corresponding to different elution volumes for all GRPELs, indicating heterogeneity in the oligomerization state of the protein (Figure S1 a-d). The major peaks, eluted at approximately 13 ml in the 24 ml SEC column, were collected and concentrated. To analyse differences between GRPEL2 and its mutants, we conducted reduced and non-reducing SDS-PAGE followed by immunoblotting. This revealed distinct bands corresponding to several molecular masses for all GRPEL2 proteins. Additionally, we observed differences in oligomerization between GRPEL2-C87A and wild type GRPEL2, but not between GRPEL2-C97A and wild type (Figure 2b). These observations imply the dynamic nature of GRPELs and their ability to exist in a multi-oligomeric state while maintaining solubility in solution.

**Figure 2.**
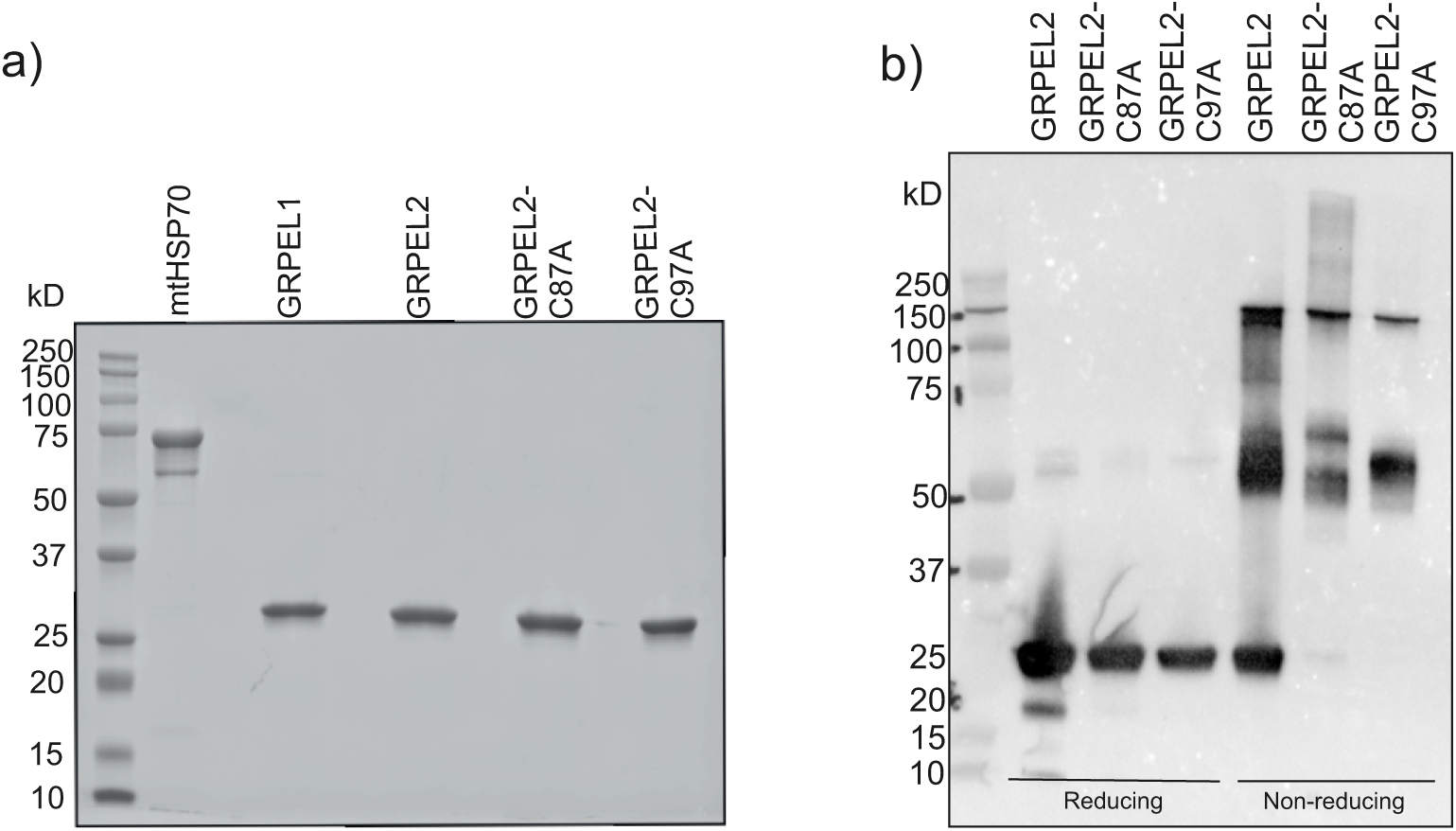
Purified-soluble human GRPELs exist in higher oligomeric forms. (a) Proteins treated with beta-mercaptoethanol and subjected to a 10-min boiling step, then run in a 10% SDS gel, show mtHSP70 at 70 kDa, and GRPEL1, GRPEL2, GRPEL2-C87A, and GRPEL2-C97A at 25 kDa. (b) Western blot of GRPEL2, GRPEL2-C87A and GRPEL2-97A, in reducing and non-reducing conditions. The non-reducing samples, GRPEL2-C87A show difference in oligomerization compared to wild type GREPEL2 and C97A, showing the effect of C87 in oligomer formation.

### 2.3 Structural stability of GRPELs depends on the interactions of cysteines

We aimed to understand the role of cystines in the stability of GRPELs using circular dichroism (CD) spectrophotometry. We compared the CD spectrums of wild type GRPEL1 and GRPEL2 with those of mutant GRPEL2-C87A and GRPEL2-C97A (Figure 3, Figure S2). The far UV CD spectra measurement at 180-280 nm indicated that the recombinant proteins were folded and soluble. In the far UV spectrum, all the recombinant GRPEL proteins were mostly α-helical (31-38%) with a smaller number of β-sheet structures (Table I). Proteins with and without DTT in the buffer were compared (Figure 3). In the absence of DTT, we noticed differences in the CD curves between GRPEL2 and GRPEL2-C87A. However, when we added DTT, these differences disappeared. At the same time, adding DTT caused notable changes in the CD spectra of GRPEL1, GRPEL2, and GRPEL2-C97A (Figure S2 a, b, and d), but there were no changes in the CD profile of GRPEL2-C87A (Figure S2 c).

**Figure 3.**
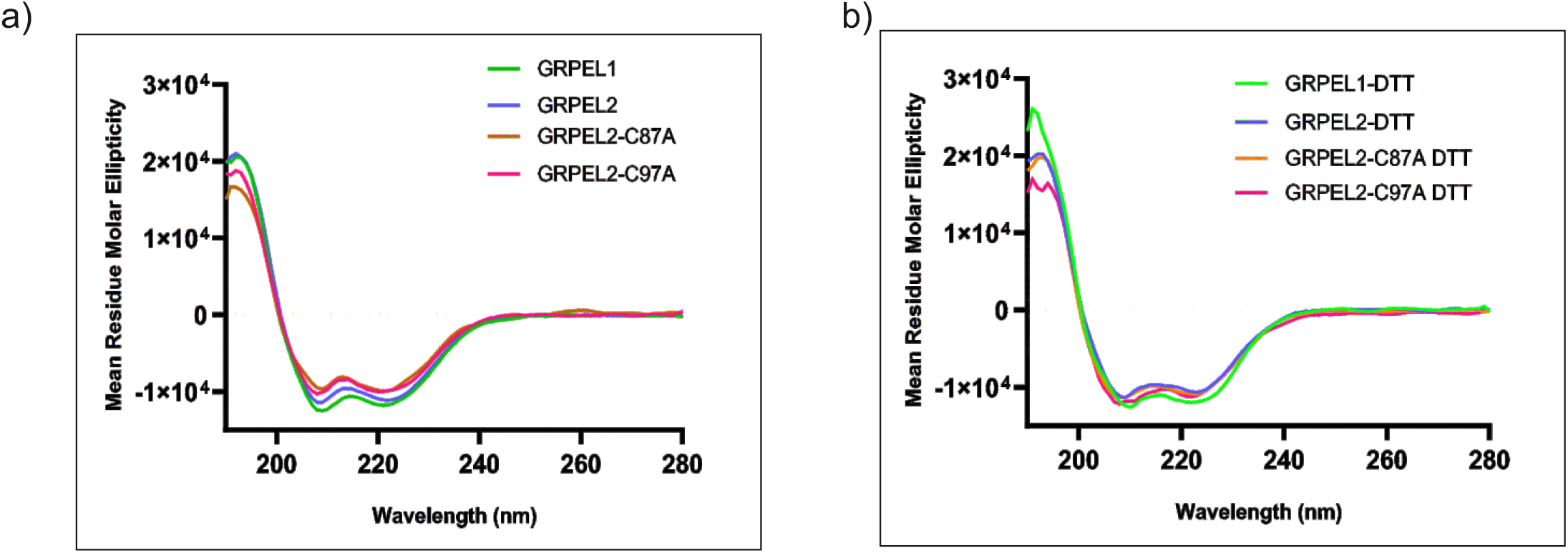
Structural stability of GRPELs depends on the interactions of cysteines. (a) GRPEL1, GRPEL2, GRPEL2-C87A, and GRPEL2-C97A represent samples diluted with buffer under nonreducing conditions. (b) GRPEL1, GRPEL2, GRPEL2-C87A, and GRPEL2-C97A represent samples diluted with buffer and the curves diluted with buffer with ImM DTT. Here, significant differences were detected with or without DTT in GRPEL1, GRPEL2, and GRPEL2-C97A but not in GRPEL2-C87A. GRPEL2 and GRPEL2-C87A have identical CD curves in the buffer with DTT, indicating the role of Cys87 in maintaining the protein conformation.

When comparing the thermal stability of proteins in buffers without DTT, GRPEL1 demonstrated higher thermal stability than GRPEL2, with a melting temperature approximately 9°C higher (Figure 4a, b). This finding aligns with previous research (Borges et al, 2003; Oliveira et al, 2006). Mutants GRPEL2-C87A and GRPEL2-C97A exhibited even higher melting temperatures, indicating greater thermo-stability compared to wild type GRPEL2 (Figure 4c, d). In the presence of DTT, GRPEL1 showed improved thermal stability compared to its state without DTT, while no marked change was observed for GRPEL2 (Figure 4e, f). Mutants GRPEL2-C87A and GRPEL2-C97A in the DTT buffer displayed stability similar to that of wild type GRPEL2 (Figure 4g, h). In conclusion, the comparison of wild type GRPEL2 with its mutants’ spectral analysis and thermal stability profiles suggests that the presence of Cys87 and Cys97 in GRPEL2 contributes to its reduced stability.

**Figure 4.**
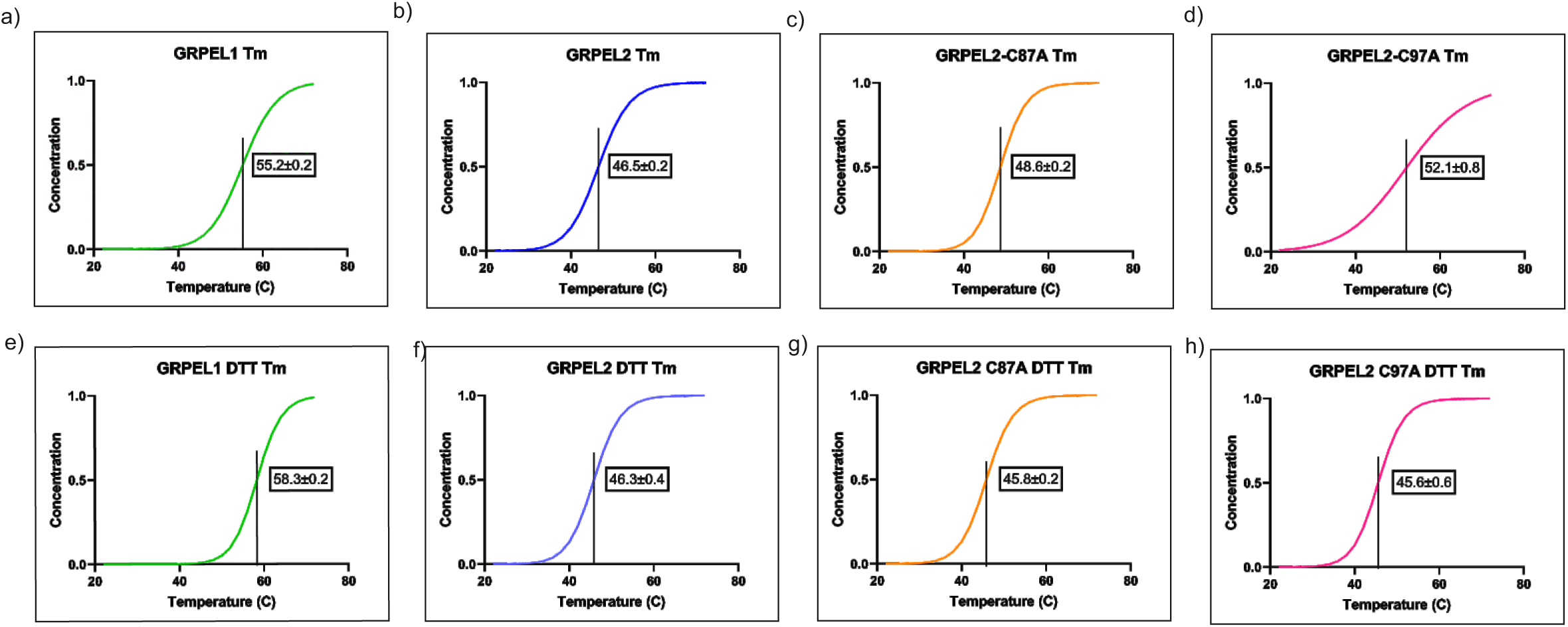
The melting temperature shows the improved stability of reduced GRPEL1. Melting temperatures in non-reducing conditions (a) GRPEL1, (b) GRPLE2, (c) GRPEL2-C87A, and (d) GRPEL2 C97A. The melting temperatures of the same proteins in the presence of reducing agents are represented in (e-h).

### 2.4 Multiangle light scattering reveals that only GRPEL1 forms a complex with mtHSP70

We characterized purified mtHSP70, GRPEL1, GRPEL2, and GRPEL2 mutants along with their possible complexes using size-exclusion chromatography coupled with multiangle light scattering (SEC-MALS). This method reliably determined the absolute molecular weight of proteins or complexes without additional standard proteins. Wild-type GRPELs displayed two distinct peaks (Figure S3), with the major peak corresponding to dimeric forms at 13ml (51 kDa for GRPEL1 and 49 kDa for GRPEL2) and the minor peak to tetrameric forms at 11.2 and 11.5 respectively (92 kDa for GRPEL1 and 87 kDa for GRPEL2) (Figure S3 a and b). Addition of 1 mM DTT reduced aggregated protein levels but did not alter GRPEL2 elution volumes (Figure S3e). Most mtHSP70 eluted as monomeric protein at 13.7 ml (70.3 kDa), with a minor fraction showing higher oligomers (166 kDa). GRPEL2-C87A and GRPEL2-C97A variants exhibited SEC profiles similar to wild type GRPEL2 (Figure S3 c and d), indicating predominantly dimeric and tetrameric forms unaffected by reducing agents, suggesting non-essential roles for cysteines in coiled-coiled region for dimerization.

The possible heterocomplexes of GRPELs with mtHSP70 were studied with SEC-MALS. GRPEL1 or GRPEL2 proteins were mixed with mtHSP70 protein in ratios of 1:1 or 3:1. In the experiments with 1:1 ratio, the SEC profiles resembled the mtHSP70 SEC profile. However, when excess GRPEL1 was used, two additional peaks appeared in the SEC profile (Figure 5a, Figure S4a and Figure S4c), indicating complex formation. The sizes of these complexes were 107 kDa (major peak, eluted in 12.2 ml) and approximately 199 kDa (minor peak, eluted in 10.6 ml). These results suggest that 2:1 and 2:2 GRPEL1 and mtHSP70 complexes were formed (expected MWs of 123 and 197 kDa, respectively). In the case of GRPEL2 or its mutant variants, no complex formation with mtHSP70 was detected (Figure 5b. Figure S4b and Figure S4d). Both the dimeric form of GRPEL2 and the monomeric form of mtHSP70 eluted at the same peak. The addition of ADP and DTT to the running buffer did not significantly alter the SEC profiles. However, the presence of DTT reduced the amount of aggregated protein. The SEC-MALS findings decisively demonstrate the exclusive formation of a visible complex between GRPEL1 and mtHSP70, starkly contrasting with the absence of such interaction with GRPEL2 or its mutants.

**Figure 5.**
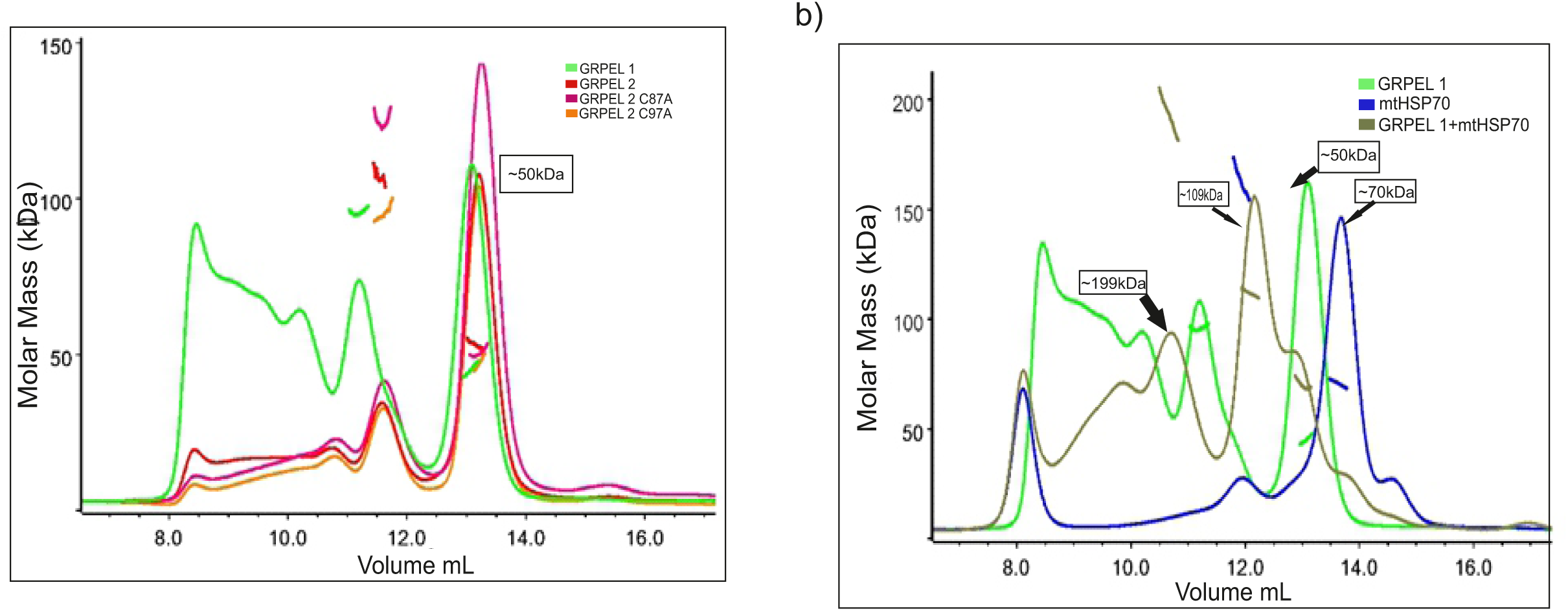
Multiangle light scattering reveals that only GRPEL1 forms a complex with ADP-bound mtHSP70. The SEC-MALS analyses of GRPELs and mtHSP70 and their potential complex. (a) The light scattering signals (angle=90°) of GRPEL1 (green), GRPEL2 (red), GRPEL2 C87A (magenta) and GRPEL2 C97A (orange) are shown (as eluted from the Superdex200 10/300GL Increase column (GE Healthcare)). In addition, the molecular mass distribution profiles of each run are shown for the peak regions (horizontal lines). The majority of each eluted GRPEL variant eluted in 13 ml as dimers with calculated molecular mass of around 50 kDa. (b) The SEC-MALS results of mtHSP70 alone (blue), GRPEL1 (green) and the complex formed between them (brown). The mtHSP70 eluted in 13.7 ml with calculated molecular mass of 70 kDa indicating monomeric nature of the protein. Interestingly, the elution volume of mtHSP70 was later than that of the dimeric GRPEL1 being smaller in the molecular weight (∼50 kDa).

### 2.5 mtHSP70 has partiality to GRPEL1 over GRPEL2 in ADP-bound state

To investigate the direct binding and affinity of the isolated proteins, we used purified mtHSP70, GRPEL1, GRPEL2, GRPEL2-C87A, and GRPEL2-C97A both in reducing and non-reducing conditions and analyzed their interaction with mtHSP70 using microscale thermophoresis (MST). In our investigation, we studied the binding of GRPEL1 and GRPEL2 with mtHSP70 under reducing conditions, both in the presence and absence of DTT and ADP. We observed that mtHSP70 bound to GRPEL1 with a K_D_ of 155 nM (Figure 6a) in the absence of ADP and 7.89 nM in its presence (Figure 6b). Conversely, GRPEL2 exhibited much lower affinity, binding to mtHSP70 with a K_D_ of 196 nM without ADP (Figure 6c) and 942 nM with ADP (Figure 6d) (Table II). Thus, the presence of ADP improved the affinity between GRPEL1 and mtHSP70 while the opposite impact was observed for GRPEL2, showing ADP binding to the chaperone regulates its NEF preference.

**Figure 6.**
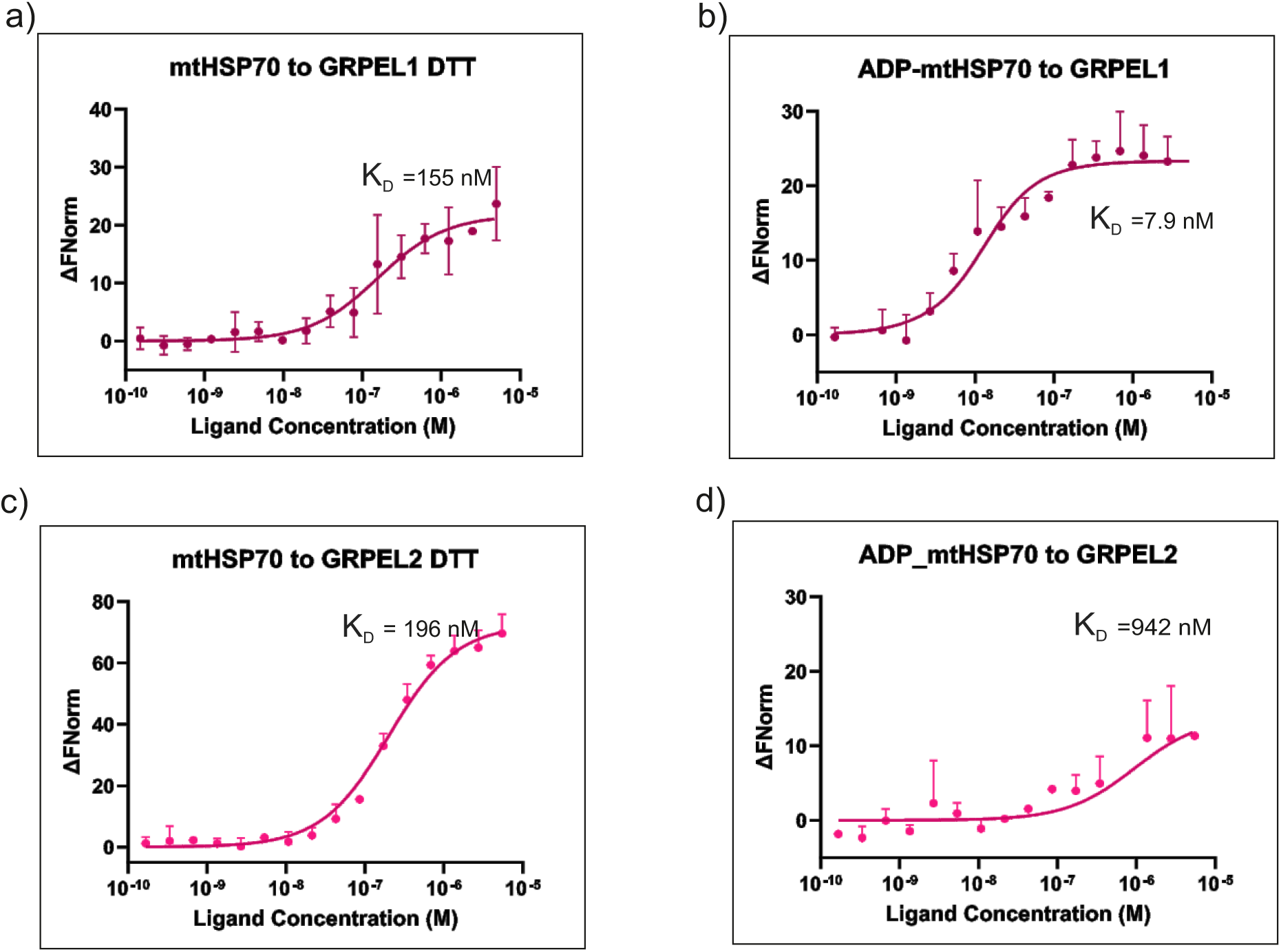
GRPEL1 predominantly binds to ADP-mtHSP70 with high binding affinity. The MST curve represents the binding of the ligand (a) GRPEL1 to mtHSP70 in presence of DTT, showing an affinity of K_D_=155nM and (b) GRPEL1 to ADP-mtHSp70, showing an affinity of K_D_=7.9nM. While c) GRPEL2 to mtHSP70 in presence of DTT (K_D_=196nM and d) GRPEL2 to ADP-mtHSP70 with affinity of K_D_=942nM Here, the binding affinity varies between both proteins, showing preference for GRPEL1 over GRPEL2.

Finally, in the non-reduced conditions, mtHSP70 bound to GRPEL1 with a K_D_ of 320 nM and GRPEL2 with a K_D_ of 520 nM (Figure S6a and Figure S5b). These results show that both GRPELs have better affinity in the reduced form in comparison with non-reduced conditions. In the same non-reducing conditions, the mutant GRPEL2-C97A bound with a K_D_ 450 nM. Interestingly, GRPEL2-C87A had a better interaction with mtHSP70 (with K_D_ 210 nM) than GRPEL1, suggesting that the disulfide bond reduces the interaction of GRPEL2 with mtHSP70 (Figure S5 c and Figure S5 d) (Table III).

### 2.6 GRPEL1 facilitates cleft opening for ADP release from the mtHSP70 nucleotide binding domain, contrasting with GRPEL2

To investigate the interaction patterns between GRPELs and mtHSP70, we utilized Alphafold-multimer to model GRPELs with mtHSP70 with 2:1 and 2:2 stoichiometry. In the resulted complexes the full-length mtHSP70 takes an unexpected conformation, which resembles the ATP-bound state of the chaperone (Figures S6 a and b). Because in this study we were mostly interested in the interactions between GRPELs and the NBD of mtHSP70, the calculations were also done accordingly by following the approach outlined by Rossi et al. (Evans et al, 2021; Jumper et al., 2021; Rossi et al., 2024) (Figure 7). The modelled complex structures suggest that both GRPELs predominantly interact with the NBD via the middle portion of the coiled-coil region (residues Lys79-Leu97) and the C-terminal β-sheet domain, including beta strands 2, 3, and 4 (see alignment in Figure 1, Figure 7a, b, Figure S6 a and b). In the complex structure of GRPEL1 and NBD, also the α-helices α2 and α3 of the four-helical bundle are forming interactions with the IIB region of NBD (Fig. 7). This interaction is critical to induce the opening of the nucleotide-binding pocket as seen in the modelled *E. coli* GrpE-NBD complex and in the cryo-EM structure of the *M.tuberculosis* GrpE-DnaK complex (Fig. 7c). This opening facilitates the release of the bound ADP molecule from NBD. In the calculations of GRPEL2 and NBD, similar interactions with the region IIB of NBD and the a-helical bundle region of GRPEL2 did not form, leaving the nucleotide-binding pocket in closed conformation (Figure 7b). This conformation resembles the GrpE-DnaK crystal structure of *E. coli*, where a point mutation (G122D) in the four-helical bundle of GrpE hindered the activation of the IIB region of NBD (Harrison et al., 1997)(Figure S7). Therefore, our modelling studies suggest that GRPEL1’s interaction with mtHSP70 NBD resembles a functional NEF interaction, while GRPEL2 modelling does not support the opening of the nucleotide-binding pocket and the following ADP release and NEF function.

**Figure 7:**
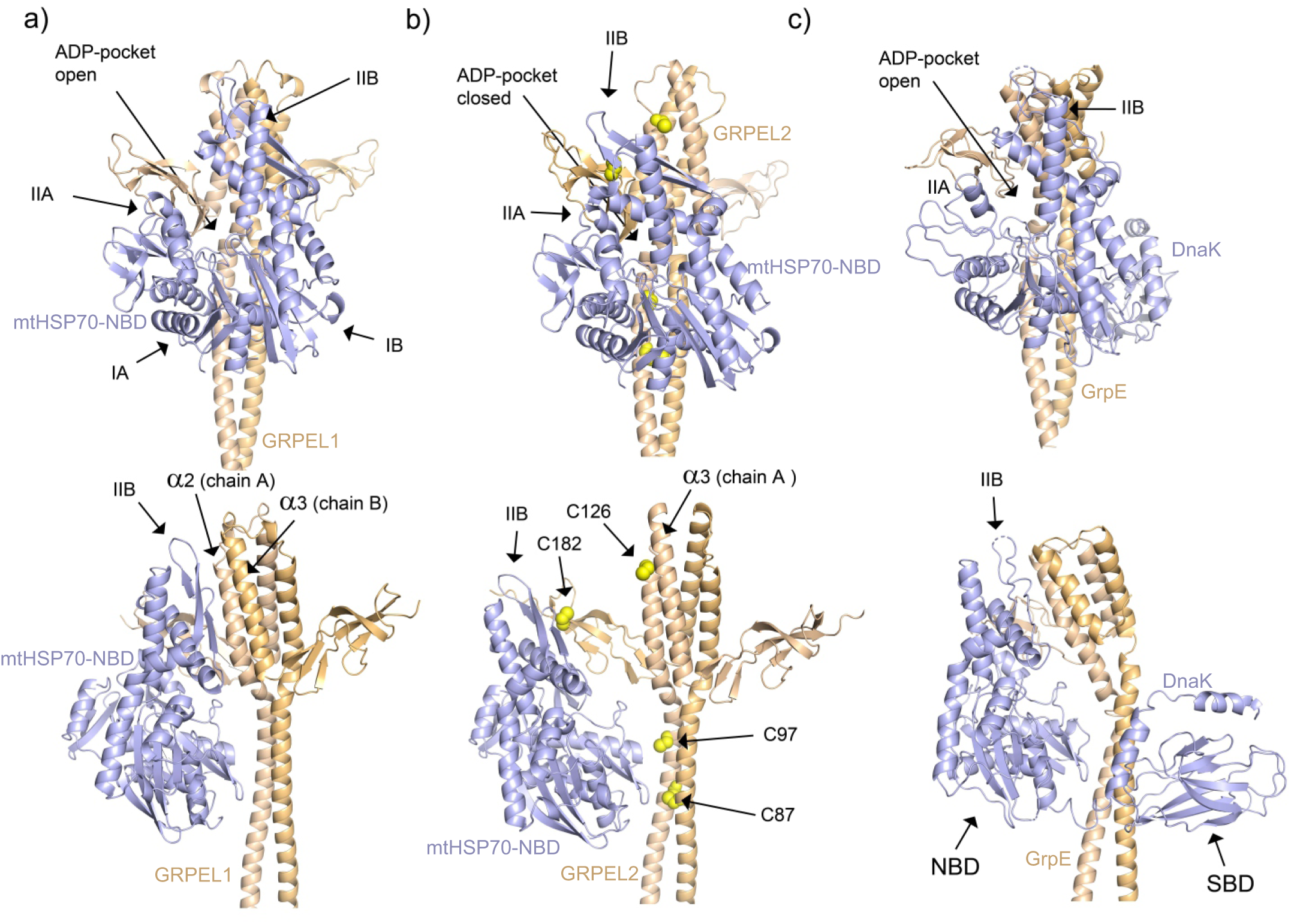
The modelling studies of GRPEL1 and GRPEL2 with the nucleotide binding domain of mtHSP70. The AlphaFold model of homodimeric GRPEL1 (a) and GRPEL2 (b) complexed with NBD of mtHSP70 shown with two different views. The top view provides a clearer depiction of the open and closed conformations of the NBD of HSP70, while the bottom view offers a more detailed representation of the GRPEL dimer and its interaction sites with the chaperone. The two chains of the GRPEL homodimers are colored light brown and orange, and the NBD of mtHSP70 in blue. In the case of GRPEL2, it also highlights the critical cysteines. (c) Cryo-EM structure of Tb GrpE (light brown and orange) and the bound DnaK (blue). The NBD and SBD are labelled. NBD contains four subdomain regions referred to as IA, IIA, IB and IIB. The nucleotide-binding pocket is formed in between subdomains IIA and IIB as shown in the figure. The four-helical bundle region, formed by helices α2 and α2, are heavily interacting with the subdomain IIB of NBD facilitating the fully open nucleotide-binding pocket in the *M. tuberculosis* GrpE-DnaK structure (panel c). This critical interaction is also seen in the modelled complex structure of GRPEL1 and NBD (panel a), whereas in the modelled complex of GRPEL2-NBD this interaction is not seen at all (panel c).

## 3 DISCUSSION

Mitochondrial protein balance plays a crucial role in cellular health and metabolism. To maintain proteostasis, it is necessary to regulate and adapt the mitochondrial protein import machinery to match the cellular requirements (Baker et al, 2007). Earlier studies by us and others have suggested that GRPEL1 is the housekeeping NEF for mtHsp70 in mitochondrial matrix protein import (Konovalova et al., 2018; Neupane et al., 2022; Srivastava et al., 2017). In contrast, loss of GRPEL2 in cultured human cells did not affect protein import, but a role for GRPEL2 in oxidative stress was proposed (Konovalova et al., 2018). Despite the prominent structural resemblance of these two paralogs, we here demonstrate the biophysical and biochemical differences between human recombinant GRPELs, which support their functional roles.

We observed that upon addition of ADP, the affinity between GRPEL1 and mtHSP70 increased approximately 20-fold, with a determined K_D_ value of 7.9 nM, which is even lower than that previously observed with bacterial GrpE and DnaK (Harrison et al., 1997; Srivastava et al., 2017). Conversely, with the addition of ADP, the affinity for GRPEL2 was nearly 100-fold lower, strongly indicating that the interaction between GRPEL1 and ADP-mtHSP70 is unequivocal and could be exclusive. In the recent cryo EM structure of *M. tuberculosis* GrpE-DnaK, the preferential stoichiometry between prokaryotic GrpE and DnaK was shown as 2:1, while the 2:2 stoichiometry may hinder the NEF flexibility for the allosteric regulation of the chaperone (Xiao et al., 2024). Our SEC-MALS analysis showed that GRPEL1 forms two complexes with mtHSP70, with stoichiometries of 2:1 and 2:2, corroborating findings from the earlier crystallographic studies with bacterial GrpE-DnaK complexes (Harrison et al., 1997; Wu et al., 2012). Yet, our modelled complex favours 2:1 interaction for the functionality of GRPEL1.

Previous studies of the *G.kaustophilus* GrpE-DnaK complex crystal structure and the *M. tuberculosis* GrpE-DnaK complex cryo-EM structure have revealed interactions between the GrpE homodimer’s four-helical bundle region and DnaK’s NBD (Wu et al., 2012; Xiao et al., 2024). This interaction, involving the β-hairpin structure near DnaK’s NBD (region IIB), aids in ADP release from DnaK. In the *E. coli* GrpE-DnaK complex, this interaction is disrupted by the G122D point mutation (Harrison et al., 1997). However, recent AlphaFold modelling suggests that wild type *E. coli* GrpE can open DnaK’s NBD through similar interactions (Rossi et al., 2024). Our AlphaFold-based modelling suggests that GRPEL1 can interact with region IIB of the NBD of mtHSP70, opening its nucleotide-binding pocket. However, in our GRPEL2-NBD modelling, this interaction was absent, leaving the NBD pocket closed. This aligns with our MST studies, showing a significant loss of affinity in the presence of ADP in the GRPEL2-mtHSP70 mixture. Additionally, SEC-MALS experiments revealed no complex formation between GRPEL2 and mtHSP70. Both GRPELs maintain a fully symmetric homodimeric structure in our calculated complexes, differing from GrpE-DnaK structures where GrpE curves towards the NBD (Wu et al., 2012; Xiao et al., 2024). We propose that the C87-C87 disulfide bridge in GRPEL2’s coiled-coil region renders its homodimer too rigid for bending, hindering interaction with mtHSP70. Differences in amino acid sequences in α-helices α2 and α3 may also reduce GRPEL2’s affinity with NBD region IIB, particularly in its ADP-bound conformation requiring coiled-coil flexibility.

We earlier showed that oxidative stress induced by hydrogen peroxide in cultured human cells increased GRPEL2 dimer formation, a phenomenon contingent upon Cys87 (Konovalova et al., 2018), a cysteine unique to GRPEL2. On the contrary, in yeast Mge1 Met155 responds to oxidative stress, enhancing monomers (Marada et al., 2013). Based on our findings in the present study, we propose that the Cys87 disulphide bridge is not critical for GRPEL2 dimerization in aqueous solution. The findings from SEC-MALS analysis validate that both the C87A and C97A mutants of GRPEL2 exhibit a behaviour alike to the wild type, predominantly existing as dimers with a minority of tetramers. This suggests that the presence or absence of Cys residues does not significantly affect the quaternary structures of the protein. Moreover, we postulate that under conditions of oxidative stress, Cys87 plays a role in stabilizing the long alpha helix within GRPEL2, potentially improving the protein’s overall functionality during stress. However, comprehensive exploration of this phenomenon is warranted to elucidate its full implications.

Disulfide bonds and protein dimerization are known to notably enhance the stability of secondary structures (Betz, 1993; Fass, 2012). The GRPELs showed a preferred oligomeric state of dimers and a low number of tetramers in the SEC-MALS analysis. These higher-order oligomeric states likely contribute to enhanced stability of the secondary structure. This was corroborated by CD spectra, confirming proper protein folding and functionality. Notably, GRPEL1 exhibited greater thermal stability compared to GRPEL2 in non-reduced buffer, consistent with previous findings (Borges et al., 2003; Oliveira et al., 2006). However, mutations in GRPEL2 (C87A and C97A) resulted in improved stability under non-reducing conditions. This can be attributed to the prevention of non-functional disulphide bonds, thus preserving the native protein structure (Karimi et al, 2016). Moreover, GRPEL1 showed enhanced thermal stability under reduced conditions, where the conformation becomes more flexible and less constrained, as suggested by earlier studies (Creighton, 1988). Overall, these observations suggest that GRPELs in reduced conformation exhibit increased stability and improved resistance to aggregation. Our MST studies and CD analysis showed that both GRPELs had better affinity under reduced conditions. The affinity between GRPEL2-C97A and mtHSP70 was closer to that of wild type GRPEL2 in the non-reduced form. Notably, the GRPEL2-C87A variant, with a K_D_ of 210 nM, was comparable with the K_D_ of 196 nM between wild type GRPEL2 and mtHSP70 in the presence of DTT. This implies that the Cys87 disulphide bond of the GRPEL2 dimer reduces GRPEL2 interaction with mtHSP70.

The remaining cysteines were not investigated in this study; nevertheless, there is a possibility that they contribute to redox sensing. Specifically, Cys126 and Cys182 warrant further exploration to understand their potential redox-regulatory roles. Our model and known GrpE-DnaK structures indicate that both cysteines reside in a region conducive to protein-protein interactions. Notably, Cys182 is exclusive to GRPEL2. These findings hint that if GRPEL2 functions as a redox sensor, it stabilises by forming disulfide bonds to an already existing dimer. It is conceivable that the rigidity of the coiled-coil helices plays a crucial role in this context.

The chaperones such as mtHSP70 are susceptible to constant allosteric changes where NEFs and the substrate protein impact chaperone’s structure (Li et al, 2016). A prior study revealed that a network mediated by Mg^2+^ ions effectively enhanced the affinity of human recombinant mtHSP70 for ADP. Interestingly, NEFs were found to systematically dismantle this network, leading to a gradual release of ADP and creating conditions conducive for ATP binding (Arakawa et al, 2011). Similarly, ADP-bound mtHSP70 may exhibit a conformational change, establishing a distinct preference solely for GRPEL1. Post-translational modifications are known to modulate the interaction dynamics of chaperones, as demonstrated by recent findings regarding the acetylation of specific lysine residues in mtHSP70 (Gao et al, 2024). Similarly, ADP may employ a crucial influence on the selection of the co-chaperone for mtHSP70 by inducing conformational changes in the chaperone protein. This intricate interplay between nucleotide binding, post-translational modifications, and co-chaperone selection highlights the sophisticated regulatory mechanisms governing the intricate protein folding processes within the mitochondrial matrix.

In summary, our study illustrates the distinct binding preferences between ADP-mtHSP70 and GRPEL1, highlighting GRPEL1 as the primary co-chaperone for mtHSP70. Conversely, the presence of disulfide bonds in GRPEL2 appears to diminish its affinity for ADP-bound mtHSP70, thereby favouring GRPEL1 for functional interactions. Although GRPEL2 predominantly adopts an oligomeric state facilitated by disulfide formation, this process imposes rigidity on the alpha helix, compromising its stability and affinity for mtHSP70. These insights shed light on the contrasting roles of GRPEL1 and GRPEL2 in human cells, providing structural clarity in their functional distinctions.

## 4 MATERIALS AND METHODS

### 4.1 Cloning and protein purification

The GRPEL constructs (contain no mitochondrial targeting sequence and presence of TEV cleavage site) encoding full-length GRPEL1, GRPEL2, GRPEL2-C87A, and GRPEL2-C97A were generated by PCR amplification using a forward primer containing the NcoI site and a reverse primer containing the Acco65I site followed by a STOP codon (TAA) (Table S2). The amplified product was digested using NcoI and Acco65I and ligated into a linear (double digested) pETHis1a (+) plasmid. The resulting sequences were verified by Sanger sequencing. The mtHSP70 was expressed from a pRSFDuet-1 plasmid coding for human mtHSP70 and yeast Zim17, and was kind gift of Professor Patrick D’Silva (Department of Biochemistry, Indian Institute of Science, Bangalore, India).

Proteins were expressed in BL21(DE3) strain and grown overnight at 37°C. Secondary cultures reached an OD of 0.6 at 30°C before induction with 0.5 mM IPTG for 12 h. Cells were collected by centrifugation at 4000g for 10 min at 4°C, then lysed in 150 mM Tris pH 8, 200 mM NaCl, 5% glycerol, 50 mM imidazole with 0.5 mg/mL lysozyme, and incubated for 45 min at 4°C followed by 0.2% DOC for 15 min at 4°C. The lysate was centrifuged at 18000g for 30 min at 4°C. Ni-NTA beads were incubated with the supernatant at 4°C for 1 h. Unbound proteins were collected in the flow-through, and elution was performed with 8 elutions of 1ml each using elution buffer: 150 mM Tris pH 8, 200 mM NaCl, 5% glycerol, and 250 mM imidazole.

### 4.2 Size Exclusion Chromatography (SEC)

Gel filtration was performed using concentrated IMAC purified proteins of 500µl in volume using a Superdex 75 GL column and Superdex-200 10/300 GL column preparative grade size exclusion chromatography column (GE Healthcare). The column was pre-equilibrated with SEC buffer composed of 50 mM Tris-HCl, 50 mM NaCl, 100 mM imidazole, and 5% glycerol at pH 8.0. The final protein solution was concentrated at approximately 5–6 mg/ml (concentration was measured using BCA) and stored in SEC buffer in 50-l aliquot volumes at −80°C.

### 4.3 SDS-PAGE

For reducing/non-reducing SDS-PAGE, sample buffer for non-reducing (250 mM Tris-HCl (pH 6.8), 10% SDS, 30% glycerol, 0.02% bromophenol blue) and reducing (Laemmli sample buffer (Bio-Rad) including 10% β-mercaptoethanol) conditions were use. Proteins for reducing SDS-PAGE were boiled at 95°C for 10 min and spun before loading. For clear native PAGE, the sample buffer was 62.5 mM Tris-HCl, pH 6.8, 25% (v/v) glycerol, 0.01% (w/v) bromophenol blue, and the running buffer was 25 mM Tris and 192 mM glycine.

### 4.4 Static light scattering (SLS)

MALS analyses of all the GRPLEs and mtHSP70 were carried out using the miniDAWN™ MALS detector (Wyatt Technologies). For these experiments, 50-100 μL of the purified protein samples (concentration ranging 2-15mg/ml) were filtered (with 0.1 μm pore size) and loaded onto a Superdex200 10/300GL column (GE Healthcare) preequilibrated with SEC buffer. Before entering the MALS detector, the samples flow through a Optilab refractive index (RI) detector (Wyatt Technologies). The RI-signal is used to measure the concentration of the protein sample. Molecular mass, polydispersity, and other analyses were carried out using the ASTRA software (Wyatt Technologies).

### 4.5 Circular Dichroism (CD) Spectroscopy

Circular dichroism (CD) spectroscopy was performed using a Chirascan CD spectrometer (Applied Photophysics, Leatherhead, UK). CD data were collected between 280 and 190 nm at 22°C using a 0.1-cm path-length quartz cuvette. CD measurements were acquired every 1 nm with 0.5 s as the integration time and repeated three times with baseline correction. Data were processed using the Chirascan Pro-Data Viewer (Applied Photophysics) and CDNN (http://www.xn--gerald-bhm-lcb.de/download/cdnn). Direct CD measurements (θ; mdeg) were converted into mean residue molar ellipticity ([θ]MR) by Pro-Data Viewer. The Tm was measured with a temperature ramping from 22° C to 72 °C at a rate of 1 °C/min. The data were analysed using Global 3 (Applied Photophysics). Purified GRPEL1, GRPEL2, GRPEL2-C87A, and GRPEL2-C97A proteins were first diluted to 2.7 mg/ml in the 1x buffer (50 mM Tris pH 8, 100 mM NaCl, 100 mM imidazole, 5% glycerol) and then 1:54 diluted in dH2O. The final solution buffer was 0.93 mM Tris pH 8, 1.85 mM NaCl, 1.85 mM imidazole, 1.9 % glycerol. All samples were treated with 1 mM DTT. The final solution buffer was 0.93 mM Tris pH 8, 1.85 mM NaCl, 1.85 mM imidazole, 1.9 % glycerol, 1 mM DTT. The final concentrations were verified by measuring the absorbance of the protein solution at 214 or 205 nm.

### 4.6 Microscale thermophoresis (MST)

MST was performed using Nanotemper Monolith NT.115. Proteins were labelled with a second-generation red amine-reactive NT-650-NHS fluorescent dye using a Nanotemper labelling kit (MO-L011). The HSP70: HIS-tag fluorescent dye concentration ratio was 1:3. Excess dye was removed by centrifugation using Amicon ultra 10k MWCO centrifugal philtres (Millipore). The actual protein concentration was determined using a NanoDrop microvolume spectrophotometer after correcting for the extinction coefficient of the proteins. Binding experiments were performed in PBS-Tween-20 buffer (10 mM PBS, 150 mM NaCl, 0.05% Tween-20, pH 8). The binding assays were performed with a fixed concentration (5 nM) of fluorescently labelled mtHSP70 and two-fold serially diluted decreasing concentrations of unlabeled GRPEL1, GRPEL2, GRPEL2-C87A, and GRPEL2-C97A (All the GRPELs’ His-tag was cleaved at TEV-cleavage site and buffer exchange to MST buffer was performed). The reaction mixture of 16 serial diluents of the ligand and labelled protein was filled into the capillary tubes (Nanotemper, MO-K022) and loaded in the Monolith NT.115 instrument accordingly. All experiments were performed at 25 °C using a medium MST power and 20% LED power. MST measurements were performed in triplicate for each experimental setup.

### 4.7 Modelling calculations

The homodimeric models of GRPEL1 and GRPEL2 were generated using an AlphaFold Colab notebook (Jumper et al., 2021).The heterocomplex models of GRPEL1 or GRPEL2 with mtHSP70 were calculated using AlphaFold-multimer (Evans et al., 2021) in the COSMIC2 cloud platform for structural biology research and education (Cianfrocco et al).

## ACKNOWLEDGEMENTS

Riitta Lehtinen, Miia Nissilä, and Jana Pennonen are acknowledged for their technical support. The facilities and expertise of the BCO Structural Biology and Molecular Biophysics core (especially Dr. Hongmin Tu), all the members of Biocenter Finland, Instruct-ERIC Centre Finland, and FINStruct are gratefully acknowledged. We thank the funding support from the Academy of Finland Centre of Excellence (MetaStem), the Sigrid Juselius Foundation (for HT), and the Orion Foundation (for PM).

## SUPPLEMENTARY FIGURE LEGENDS

**Figure S1:**
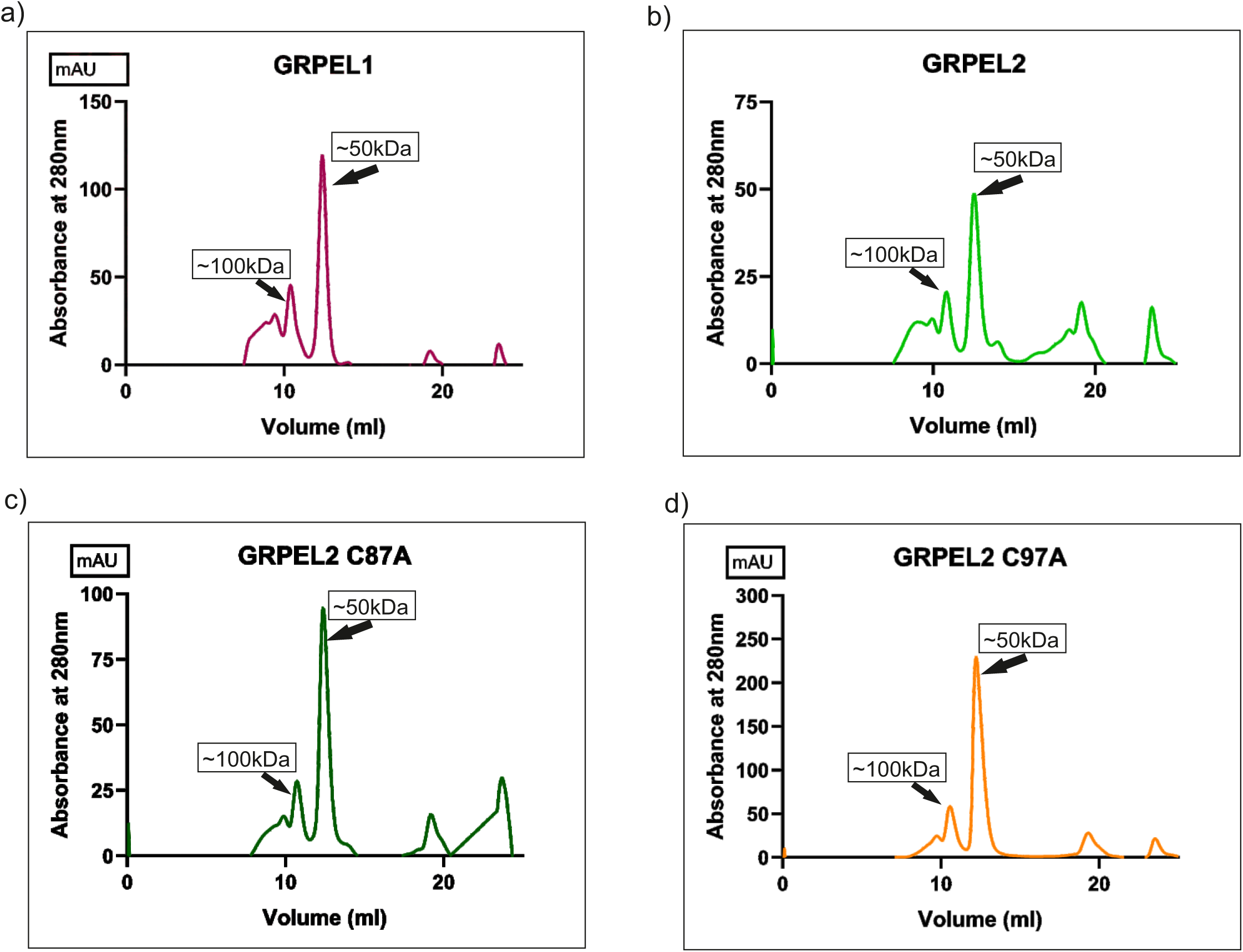
SEC chromatograms of (a) GRPEL1, (b) GRPEL2, (c) GRPEL2 C87A, (d) GRPEL2 C97A from 24ml S200 increase column used in AKTA purification system. The proteins eluted are without DTT and one major peak around volume 13ml and minor peak volume 11ml can be observed.

**Figure S2:**
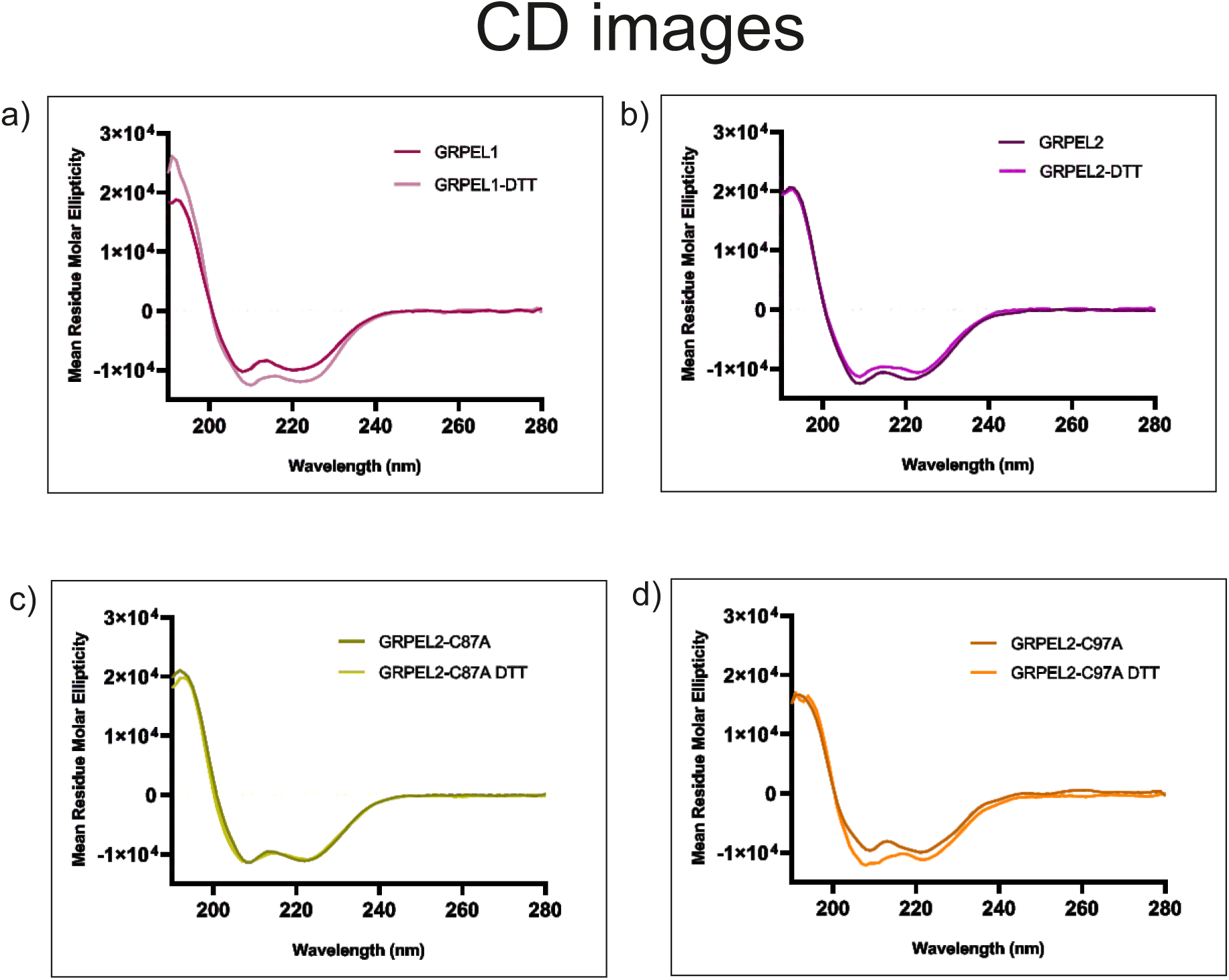
SEC chromatograms of (a) GRPEL1, (b) GRPEL2, (c) GRPEL2 C87A, (d) GRPEL2 C97A, and (e) GRPEL2 in the presence of 1 mM DTT. Shown are the UV curves at 280 nm (A280) measured by the SPD-M20A diode array detector of the high-performance-liquid-chromatography system (Shimadzu Corp.) and drawn by the Astra software (Wyatt technologies). The X-axis shows the SEC elution volume in mL and the Y axis shows the relative UV absorbance. The experiments were done in the SEC buffer 50 mM Tris, pH8, 150 mM NaCl, 5% glycerol. The addition of 1 mM DTT in the SEC buffer (panel E) did not change the elution volume indicating that any disulfide is crucial for the dimeric interactions.

**Figure S3:**
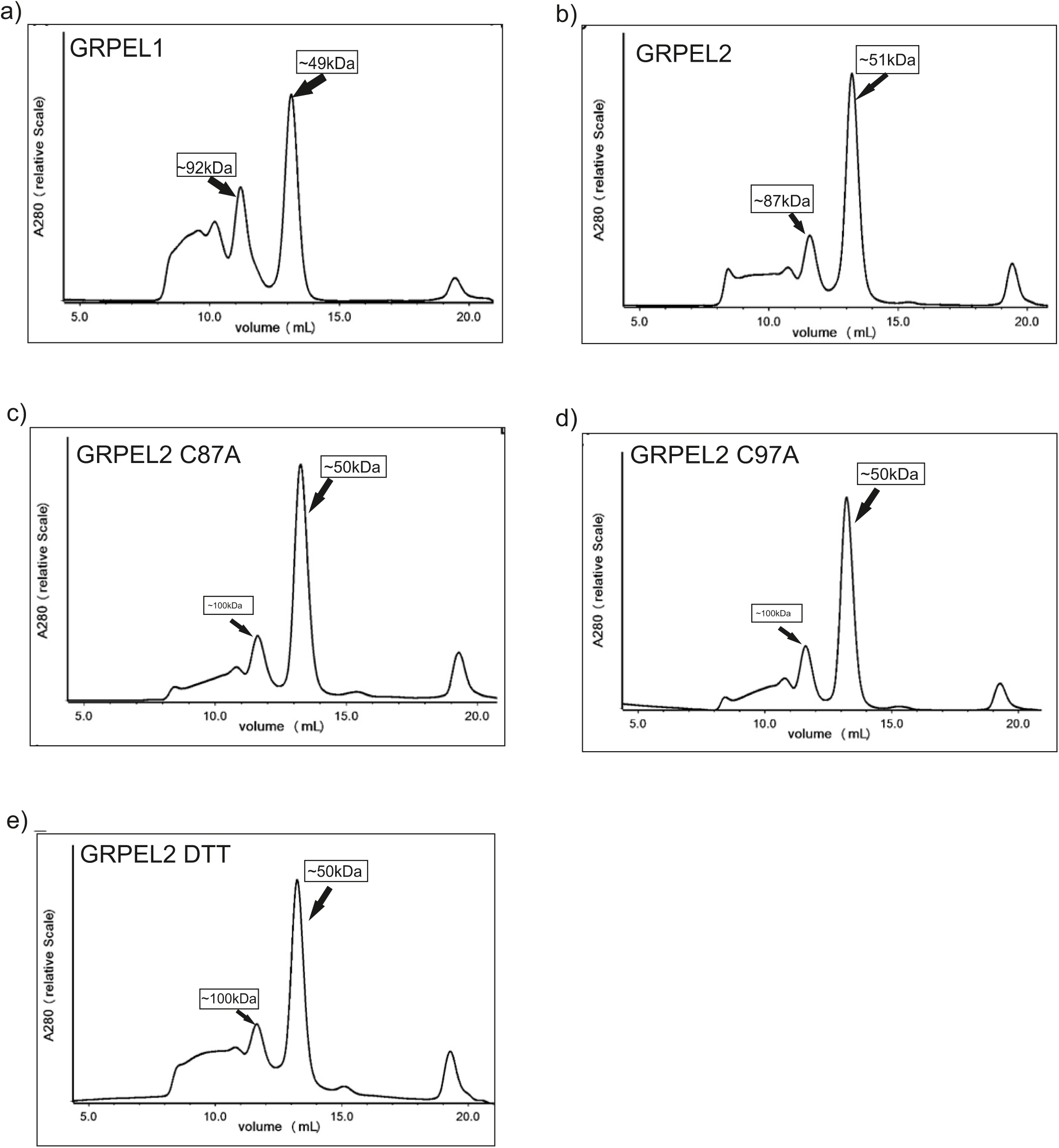
Profiles of CD runs depicting the ellipticity of protein samples in the presence or absence of a reducing agent. Panels a, b, c, and d, display the CD and Tm run profiles for GRPEL1, GRPEL2, GRPEL2-C87A, and GRPEL2-C97A, respectively.

**Figure S4:**
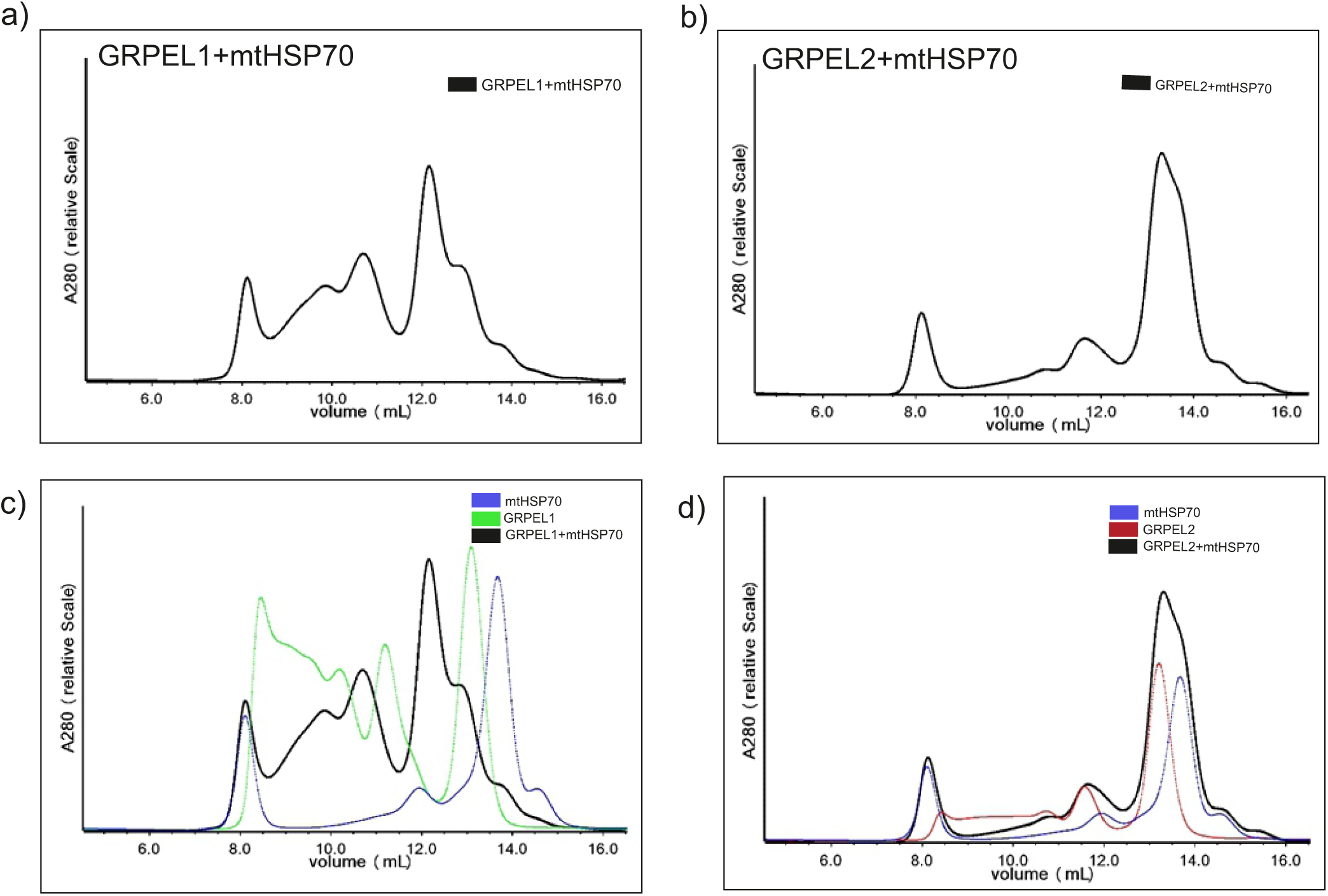
SEC chromatograms after mixing of (a) GRPEL1 and mtHSP70 and (b) GRPEL2 and mtHSP70. Shown are the UV curves (A280) measured by the SPD-M20A diode array detector of the high-performance-liquid-chromatography system (Shimadzu Corp.) and drawn by the Astra software (Wyatt technologies). The X-axis shows the SEC elution volume in mL and the Y axis shows the relative UV absorbance. (c) The GRPEL1-mtHSP70 mixture (black) led to the new peaks with shorter elution time and higher molecular weights as compared to the SEC profiles of individual GRPEL1 (green) and mtHSP70 (blue). (d) The GRPEL2-mtHSP70 mixture (black) led to one large peak which superposes perfectly with the individual SEC profiles of GRPEL2 (red) and mtHSP70 (blue) peaks. The complex of only GRPEL1-mtHSP70 can be observed while no peak shift in GRPEL2-mtHSP70 complex.

**Figure S5:**
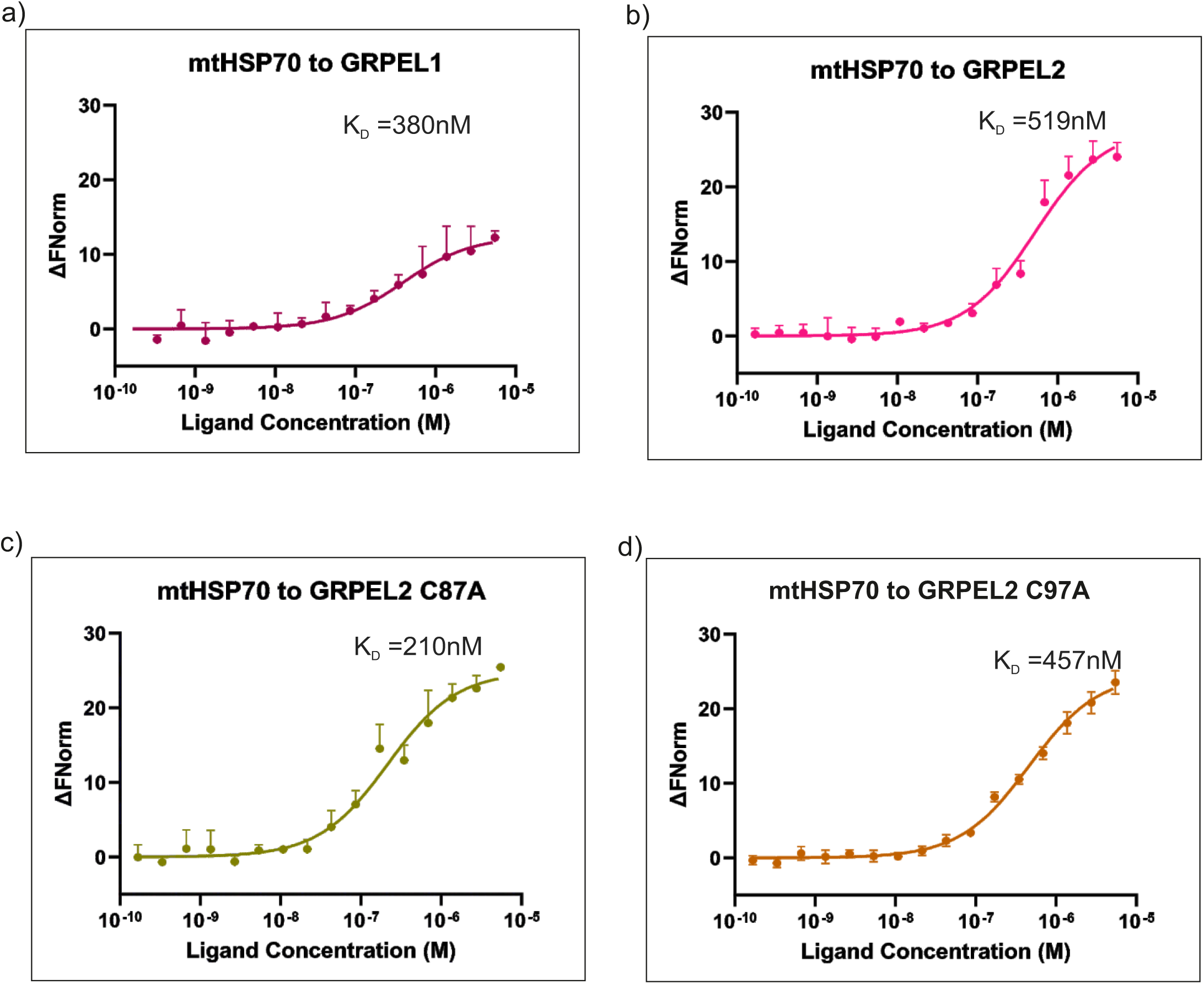
Microscale thermophoresis experiments were conducted to assess the binding affinity between the target mt-HSP70 and the ligands: (a) GRPEL1, (b) GRPEL2, (c) GRPEL2-C87A, and (d) GRPEL2-C97A. The results indicate that GRPEL1 exhibits a higher affinity than both wild-type GRPEL2 and mutant GRPEL2-C97A. Notably, mutant GRPEL2-C87A demonstrates superior affinity than all other GRPELs under non-reduced conditions.

**Figure S6.**
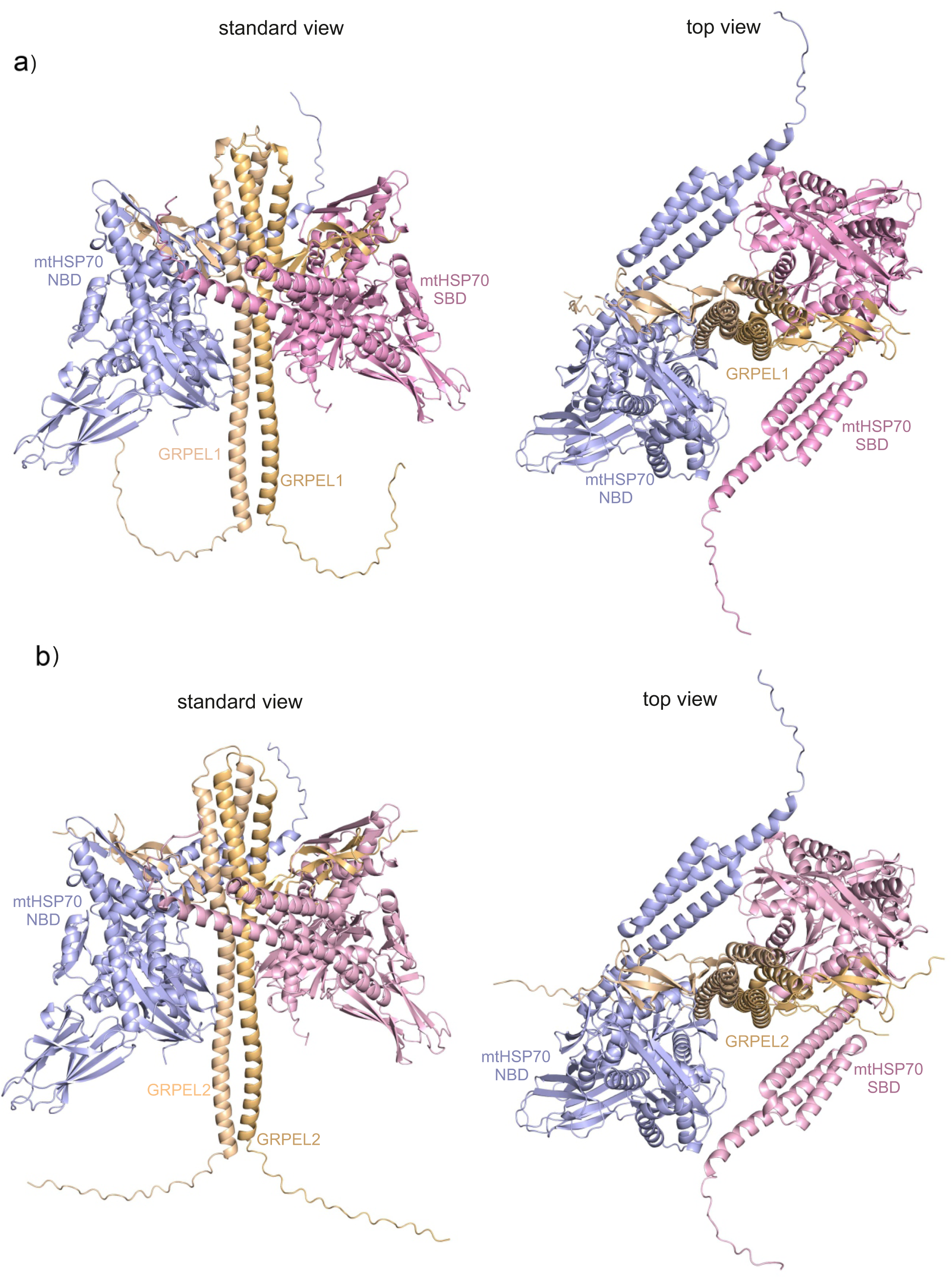
The predicted structure of the heterotetrameric complex structure of human GRPELs and mtHSP70. The AlphaFold model of homodimeric GRPEL1 (a) and GRPEL2 (b) complexed with full-length of mtHSP70 shown with two different views. The left view is the “standard view” where the central GRPEL homodimer is viewed from side (as also seen in Figure 1b and c. The right view is from the top along the GRPEL coiled-coil helices. The GRPELs are coloured in light brown (chain A) and light orange (chain B), and the mtHSP70 molecules with light blue (chain C) and pink (chain D). The subdomains of mtHSP70 are highlighted in the panel a.

**Figure S7:**
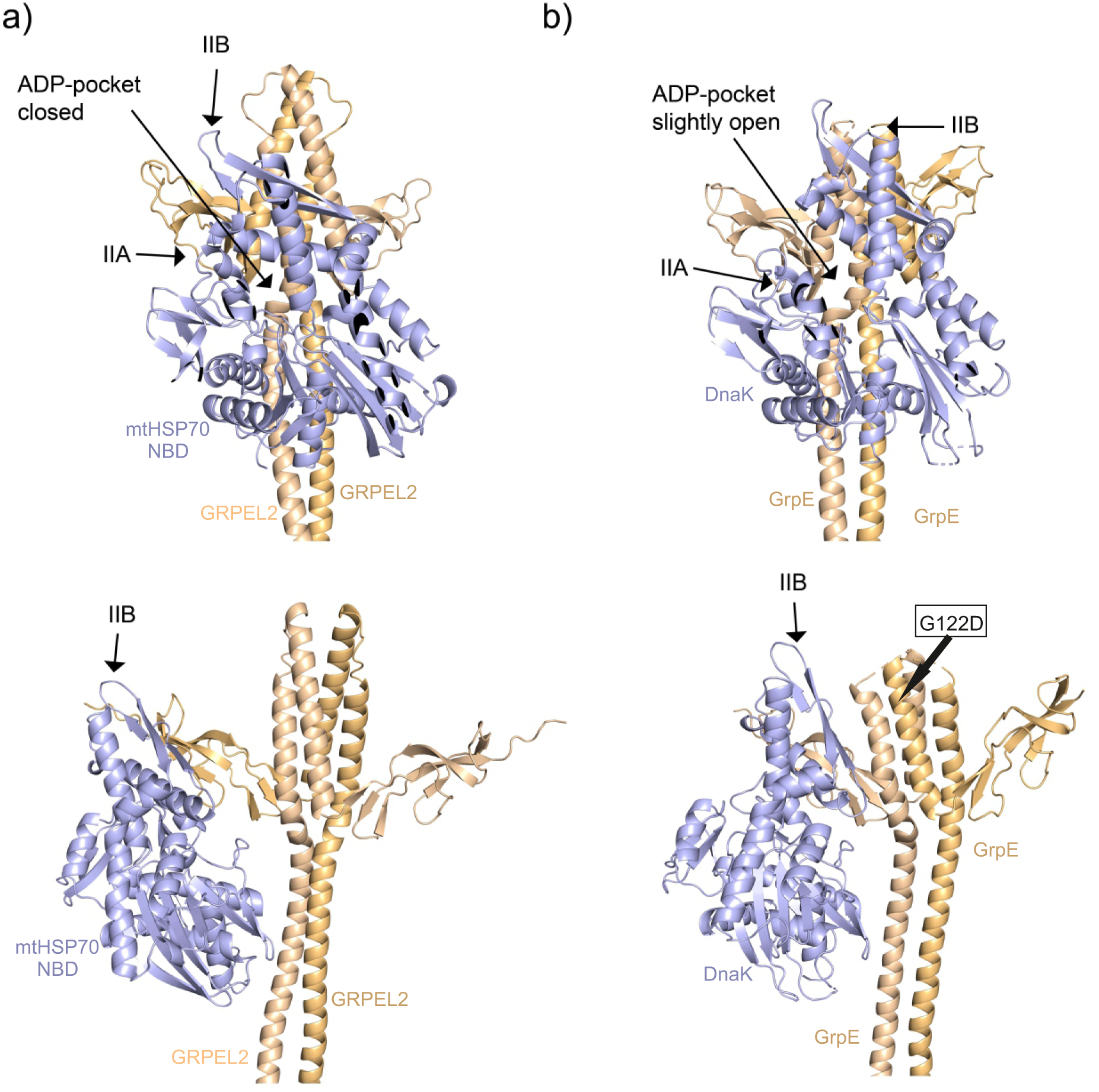
The comparison of modelled GRPEL2-NBD complex with the crystal structure of *E. coli* GrpE(G122D)-DnaK complex. a) The AlphaFold model of homodimeric GRPEL2 complexed with NBD of mtHSP70 shown in two views as in Figure 7b. b) The crystal structure of the *E. coli* GrpE-G122D in complex with the NBD of DnaK (PDB 1DKG) shown in the same views as the GRPEL2-NBD complex in panel a. The top view provides a clearer depiction of the open and closed conformations of the NBD of HSP70/DnaK, while the bottom view is a “standard view” offering a more detailed representation of the GRPEL dimer and its interaction sites with the chaperon. The two chains of the GRPEL homodimers are coloured light brown and orange, and the NBD (nucleotide-binding-domain) of mtHSP70 in blue. The subdomains IIA and IIB lining the nucleotide-binding (ADP-)pocket of NBD are labelled. The four-helical bundle region of GRPEL2 is not interacting with the subdomain IIB of NBD at all and the nucleotide-binding pocket is fully closed. In GrpE(G122D)-DnaK complex, the four-helical bundle region of GrpE is weakly interacting with the IIB subdomain of DnaK, but not enough to fully open the NBD binding pocket (as seen in the structure of M. tuberculosis GrpE-DnaK complex, Figure 7c).

---

**Table I:**
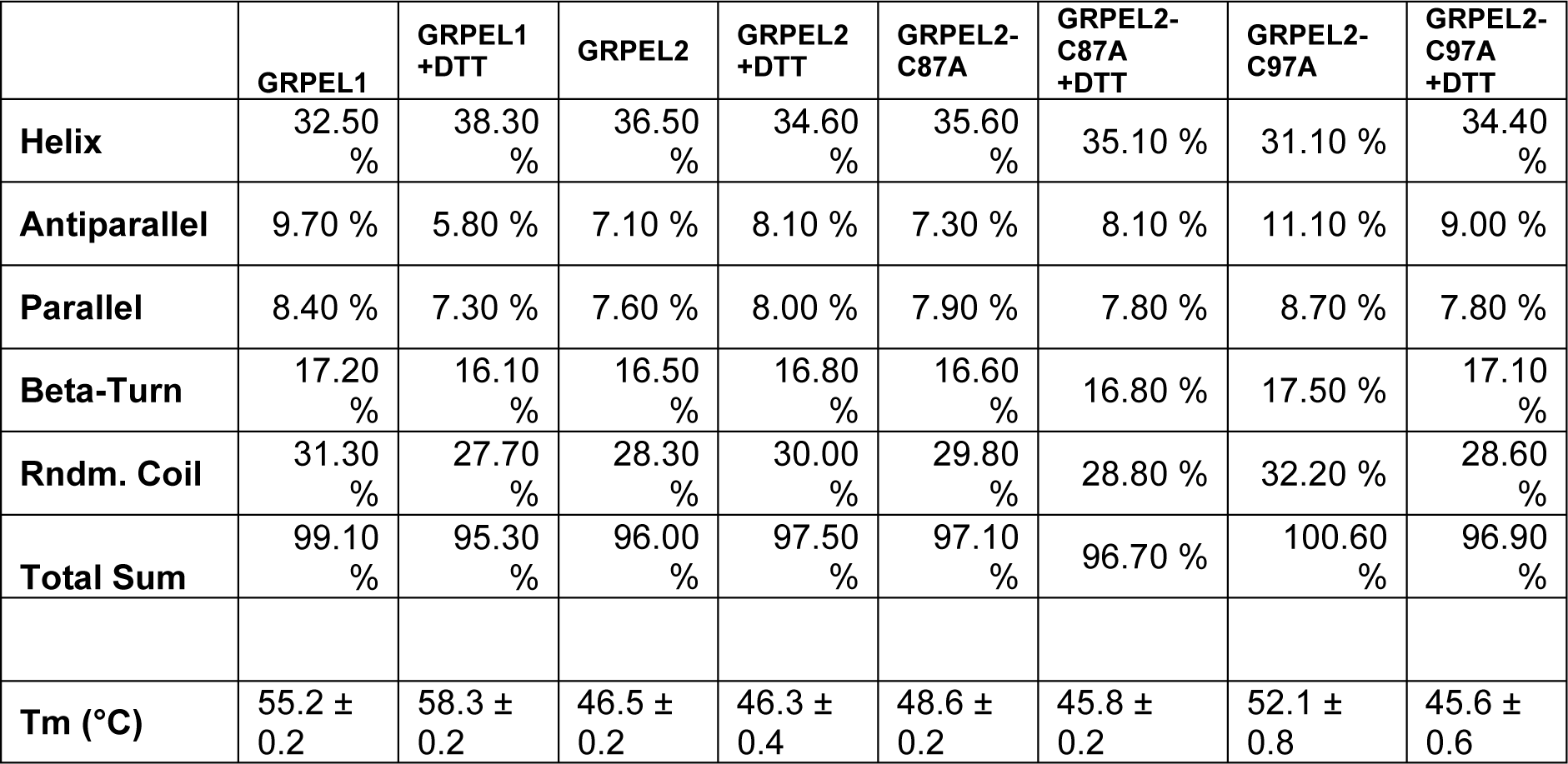
CD measurements and Tm values for various GRPELs in both the non-reducing buffer and a buffer containing DTT. Comparing Tm between GRPEL1 and GRPEL1 DTT shows increased stability in reduced proteins. Conversely, GRPEL2 samples display no significant change in Tm values. Mutants GRPEL2 C87A and C97A exhibit higher Tm values than the wild type; however, all reduced mutants show similar Tm values to wild-type GRPEL2. All GRPELs display a higher percentage of alpha helix, followed by beta sheets.

|  | GRPEL1 | GRPEL1 +DTT | GRPEL2 | GRPEL2 +DTT | GRPEL2-C87A | GRPEL2-C87A +DTT | GRPEL2-C97A | GRPEL2-C97A +DTT |
| --- | --- | --- | --- | --- | --- | --- | --- | --- |
| **Helix** | 32.50 % | 38.30 % | 36.50 % | 34.60 % | 35.60 % | 35.10 % | 31.10 % | 34.40 % |
| **Antiparallel** | 9.70 % | 5.80 % | 7.10 % | 8.10 % | 7.30 % | 8.10 % | 11.10 % | 9.00 % |
| **Parallel** | 8.40 % | 7.30 % | 7.60 % | 8.00 % | 7.90 % | 7.80 % | 8.70 % | 7.80 % |
| **Beta-Turn** | 17.20 % | 16.10 % | 16.50 % | 16.80 % | 16.60 % | 16.80 % | 17.50 % | 17.10 % |
| **Rndm. Coil** | 31.30 % | 27.70 % | 28.30 % | 30.00 % | 29.80 % | 28.80 % | 32.20 % | 28.60 % |
| **Total Sum** | 99.10 % | 95.30 % | 96.00 % | 97.50 % | 97.10 % | 96.70 % | 100.60 % | 96.90 % |
| **Tm (°C)** | 55.2 ± 0.2 | 58.3 ± 0.2 | 46.5 ± 0.2 | 46.3 ± 0.4 | 48.6 ± 0.2 | 45.8 ± 0.2 | 52.1 ± 0.8 | 45.6 ± 0.6 |

**Table II:** MST Fit Information for the interaction between mtHSP70 (target) with or without ADP bound and the ligands GRPEL1 and GRPEL2 under reducing conditions. The results demonstrate a significant difference in affinity between GRPEL1 and GRPEL2 with ADP-bound mtHSP70, whereas there is no change in affinity between GRPELs and mtHSP70 in the absence of AD

| **ADP** | **+** | **+** | **-** | **-** |
| --- | --- | --- | --- | --- |
|  | **GRPEL1** | **GRPEL2** | **GRPEL1** | **GRPEL2** |
| **Bound** | 923.27 | 907.31 | 910.30 | 947.42 |
| **Unbound** | 899.92 | 893.35 | 888.52 | 874.85 |
| **Kd** | **7.6 nM** | **942 nM** | **155 nM** | **196nM** |
| **Target Conc** | 0.00000001 | 0.00000001 | 0.000000001 | 0.00000001 |
| **Standard Deviation:** | 1.99 | 2.45 | 1.53 | 2.19 |
| **Kd Confidence:** | ±2.67103E-09 | ±7.43327E-07 | ±3.54328E-08 | ±2.1727E-08 |

**Table III:**
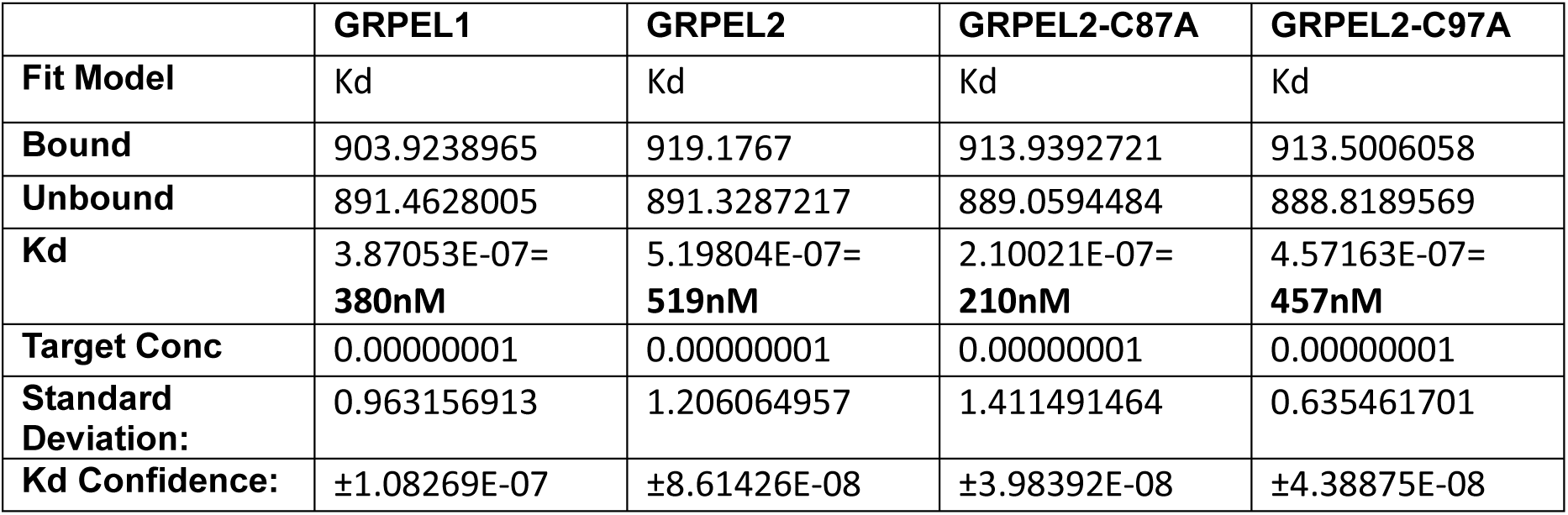
MST fit information for the interaction of mtHSP70 (target) with the ligands GRPEL1, GRPEL2, GRPEL2-C87A, and GRPEL2-C97A. In non-reducing conditions, GRPEL1 exhibits superior affinity to mtHSP70 compared with GRPEL2 and GRPEL2-C97A. Notably, GRPEL2-C87A demonstrates the highest affinity among all ligands in this context.

